# The impact of PNPLA3 and TM6SF2 in cirrhosis related complications

**DOI:** 10.1101/2021.01.07.425655

**Authors:** Xue Shao, Haruki Uojima, Taeang Arai, Yuji Ogawa, Toru Setsu, Masanori Atsukawa, Yoshihiro Furuichi, Yoshitaka Arase, Kazue Horio, Hisashi Hidaka, Takahide Nakazawa, Makoto Kako, Tatehiro Kagawa, Katsuhiko Iwakiri, Atsushi Nakajima, Shuji Terai, Yasuhito Tanaka, Wasaburo Koizumi

## Abstract

Patatin-like phospholipase domain-containing 3 (PNPLA3) and transmembrane 6-superfamily member 2 (TM6SF2) polymorphisms have major impact for fibrosis due to steatohepatitis. However, there are scant data about correlations between cirrhosis-related complications and the polymorphisms of these genes. Therefore, we aimed to determine the role of the PNPLA3 and TM6SF2 polymorphisms in fibrosis progression for patients with liver cirrhosis. A multicenter study was performed at six hospitals in Japan enrolling 400 patients with liver cirrhosis caused by virus (n = 157), alcohol (n = 104), nonalcoholic fatty liver disease (NAFLD) (n = 106), or autoimmune disease (n = 33). These cirrhotic patients included those with complications of variceal bleeding, hepatic ascites, and/or hepatic encephalopathy and those without. To assess the role of the PNPLA3 and TM6SF2 polymorphisms in patients with cirrhosis related complications, we calculated the odds ratio and relative risk for the rs738409 and rs58542926 polymorphisms. We also accessed whether or not the interaction between these two polymorphisms contributed to cirrhosis related complications. As a result, the odds ratio for complications in the NAFLD group significantly increased in the presence of the rs738409 GG genotype when the CC genotype was used as the reference. There were no significant risks between complications and the presence of the rs738409 G allele in the virus or alcohol groups. There were no significant risks of complications in the frequency of the rs58542926 T polymorphism regardless of the etiology of liver cirrhosis. The interaction between the rs738409 and rs58542926 polymorphisms had the highest odds ratio of 2.415 for complications in the rs738409 GG + rs58542926 (CT+TT) group when rs738409 (CC+CG) + TM6SF2 CC was used as the reference in the NAFLD group although there was no statistically interaction between these gene polymorphisms (P=0.085).

## Introduction

Chronic liver damage is often caused by the long-term progression of hepatic steatosis, with a substantial risk of progression to advanced fibrosis and liver-related mortality [1, 2]. The molecular mechanism of steatohepatitis remains unclear. However, many studies suggested that the accumulation of triglycerides causes lipotoxicity, which promotes cell damage induced by excessive oxidative stress [3]. Damaged hepatocytes release damage-endogenous-associated molecular patterns and directly cause hepatic stellate cell activation, which leads to the accumulation of extracellular matrix proteins and subsequently to fibrosis progression [4].

Patients with lipotoxicity-induced hepatic damage typically present with insulin resistance that is associated with type 2 diabetes, dyslipidemia, hypertension, and cardiovascular disease [5]. Among others, gene polymorphism has been assessed as a risk factor of developing steatohepatitis. Previously, the patatin-like phospholipase domain-containing 3 (PNPLA3) and transmembrane 6-superfamily member 2 (TM6SF2) genes were identified as genetic factors for steatohepatitis by the GWAS (Genome Wide Association Study) [6]. Many researchers showed that PNPLA3 and TM6SF2 polymorphisms were associated with the severity of fibrosis in patients with nonalcoholic fatty liver disease (NAFLD) and alcoholic liver disease (ALD) [7–13]. Furthermore, the PNPLA3 and TM6SF2 polymorphisms impact steatosis and liver damage in chronic virus hepatitis [14, 15].

Thus, the PNPLA3 and TM6SF2 polymorphisms have major impacts on fibrosis due to steatohepatitis. Therefore, polymorphisms of these genes should be analyzed for patients with chronic liver diseases regardless of etiology. Furthermore, different stages of liver fibrosis due to liver steatosis ultimately lead to the development of liver cirrhosis (LC) [16]. Moreover, the presence or absence of cirrhosis-related complications are the most important factors for predicting prognosis and determining the treatment strategy for patients with chronic liver disease [17, 18]. Therefore, the polymorphisms of these genes should be analyzed regardless of the stages of fibrosis. However, previously studies enrolled non-cirrhotic patients; therefore, there are scant data on the PNPLA3 and TM6SF2 polymorphisms in cirrhotic patients. Moreover, to our knowledge, there are no data about any correlations between cirrhosis-related complications and the polymorphisms of these genes. In the present study, we aimed to determine the role of the PNPLA3 and TM6SF2 polymorphisms in fibrosis progression for patients with LC.

## Materials and Methods

### Ethics

The research protocol for this study conformed to the provisions of the Declaration of Helsinki, was approved by the Institutional Review Board Ethics Committee of the Kitasato University School of Medicine (Number: G19-06), Yokohama City University Graduate School of Medicine (Number: A200528001), Nippon Medical School Chiba Hokusoh Hospital (approval number: 603) and the Tokushukai Medical Group (Number: TGE01428-024). This is registered in the UMIN Clinical Trials Registry as UMIN000039573. All patients were informed about the significance of gene analysis and gave written informed consent.

### Population

This multicenter study was performed at six hospitals in Japan. In this study, A total of 400 cirrhotic patients were assessed from November 2019 through May 2020. Patients with LC were recruited on the basis of laboratory results and imaging tests, showing an irregular surface of the liver, collateral portosystemic venous channels, ascites, hepatosplenomegaly, and/or esophageal (gastric) varices. The patients were categorized according to the presence or absence of the following life-threatening complications associated with LC: hepatic ascites, hepatic encephalopathy, and/or varix ruptures [19]. A histological evaluation by liver biopsy was not performed because of the incidence of adverse events and sampling error staging inaccuracies. LC in these patients was caused by viral infections due to hepatitis B and C viruses, ALD, NAFLD, autoimmune disease (AID). including primary biliary cholangitis, primary sclerosing cholangitis, and autoimmune hepatitis. Hepatitis B virus infection was diagnosed on the basis of the presence of the hepatitis B surface antigen and/or polymerase chain reaction (PCR) assays performed to directly detect the hepatitis B virus DNA. Hepatitis C virus (HCV) infection was diagnosed on the basis of the presence of HCV-ab and/or HCV RNA (1.2 log IU/mL or lower). ALD was diagnosed on the basis of an ethanol intake of about 40–80 g/day (men) or 20–40 g/day (women) for longer than 10 years and exclusion of other liver diseases. A definitive diagnosis of NAFLD required the presence of specific histological features. However, a liver biopsy was not performed because liver fibrosis progression lead to a reduction in hepatic fat to the point of complete fat loss. Therefore, patients with an intake of <20 g of ethanol per day, and the appropriate exclusion of other liver diseases, were categorized as those with NAFLD. Patients with severe kidney or heart disease or malignancies other than heptaocellular carcinoma were excluded.

### Genotyping for the polymorphisms in PNPLA3 and TM6SF2

Blood samples were exclusively collected in EDTA (ethylenediaminetetraacetic acid) tubes, and genomic DNA extraction was performed using an automated MagNA Pure Compact Instrument (Roche Diagnostics, Mannheim, Germany) with the MagNA Pure Compact Nucleic Acid Isolation Kit I (Roche Diagnostics) according to the manufacturer’s instructions.

Genotyping for the rs738409 and rs58542926 polymorphisms were performed using TaqMan SNP Genotyping Assays (Applied Biosystems, Waltham, USA) and LightCycler 96 System (Roche Diagnostics). The reaction consisted of 10 *µ*L LightCycler FastStart Essential DNA Probes Master Mix (Roche Diagnostics), 1 *µ*L of the primer-probe mix (20 ×), 7 *µ*L of water (PCR grade), and 2 *µ*L of the DNA sample to obtain a total reaction volume of 20 *µ*L. Genotypes of the rs738409 polymorphism were categorized as the CC genotype (wild type), CG genotype (heterozygous polymorphism), and GG genotype (homozygous polymorphism). Genotyping for rs58542926 polymorphism was categorized as the CC genotype (wild type), the CT genotype (heterozygous polymorphism) and the TT genotype (homozygous polymorphism).

### Baseline characteristics

Demographic parameters, concomitant medication, and baseline medical data was obtained on the same day as genotyping or within 1 month. Body mass index (BMI) was measured, and obesity was defined as a BMI of >25.0 kg/m^2^. Diabetes was diagnosed on the basis of the hemoglobin A1c level.

### Endpoints

To assess the role of the PNPLA3 and TM6SF2 polymorphisms in patients with LC, expression frequencies of the rs738409 and rs58542926 polymorphisms were analyzed in the enrolled patients. Subsequently, the odds ratios and relative risks for the rs738409 and rs58542926 polymorphisms were calculated for the patients with and those without complications. We also accessed whether or not the interaction between the rs738409 and rs58542926 polymorphisms contributed to fibrosis progression.

### Statistical analyses

Based on previous studies [8], the sample size was calculated to be approximately 400 patients (200 patients each with and without complications) for this study with a confidence interval (CI) of 95% and a power of 80%. Data were expressed as the mean*±*standard deviation. Genotype distribution was examined using the chi-square test.Patients with and without complications were compared using the unpaired t-test. Frequencies of the rs738409 and rs58542926 polymorphisms were analyzed using the Hardy-Weinberg equilibrium. The association between genotype and categorical variables was determined using logistic regression models. Differences with a P value of <0.05 were considered statistically significant. Data were analyzed using the SPSS version 24.0 software package (IBM Corporation, Armonk, NY, USA). Statistical analyses were performed by the Statista Corporation, Kyoto, Japan.

## Results

### Analysis and comparison of baseline characteristics

Table 1 summarizes thepatients’ demographics and other baseline characteristics. The LC in these patients was caused by virus (n = 157), alcohol (n = 104), NAFLD (n= 106), and AID (n = 33). A total of 199 (49.8%) patients had complications. The numbers of hepatic encephalopathy, hepatic ascites, and/or varix rupture were 108, 144, and 47, respectively.

**Table 1.** Patient baseline clinical characteristics

| Characteristics |  | Virus (HCV/HBV) | Alcohol | NAFLD | AID<br>(AIH/PBC) | Total |
| --- | --- | --- | --- | --- | --- | --- |
| N |  | 157 (110/47) | 104 | 106 | 33 (14/19) | 400 |
| Age | yrs | 69.2 ± 10.3 | 63.1 ± 11.6 | 67.8 ± 12.5 | 71.1 ± 8.0 | 67.4 ± 11.4 |
| Gender: Male | n (%) | 91 (58.0) | 81 (77.9) | 54 (50.9) | 7 (21.2) | 233 (58.3) |
| BMI | kg/m<sup>2</sup> | 23.4 ± 4.07 | 22.9 ± 3.26 | 26.2 ± 5.06 | 22.4 ± 3.53 | 23.9 ± 4.35 |
| Obesity | n (%) | 45 (28.8) | 27 (26.0) | 63 (59.4) | 6 (18.2) | 141 (35.3) |
| Liver cancer | n (%) | 73 (46.5) | 19 (18.3) | 20 (18.9) | 3 (9.1) | 115 (28.7) |
| HCV SVR/Non-SVR | n (%) | 79/27 | — | — | — | — |
| DM | n (%) | 50 (31.8) | 37 (35.6) | 58 (54.7) | 4 (12.1) | 149 (37.3) |
| Complication<br>HE/HA/VR | n (%) | 75 (47.8) 30/55/11 | 57 (54.8)<br>39/40/19 | 51 (48.1)<br>32/41/11 | 16 (48.5) 7/8/6 | 199 (49.8)<br>108/144/47 |
| Child-Pugh score |  | 6.03 ± 1.81 | 6.73 ± 2.10 | 6.35 ± 2.05 | 6.24 ± 1.93 | 6.31 ± 1.98 |
| FIV-4 index |  | 5.53 ± 3.69 | 5.91 ± 5.23 | 5.96 ± 4.23 | 5.87 ± 3.12 | 5.77 ± 4.23 |
| ALBI score |  | −2.39 ± 0.66 | −2.16 ± 0.75 | −2.21 ± 0.68 | −2.10 ± 0.62 | −2.35 ± 0.69 |

BMI, Body mass index ; NAFLD, nonalcoholic fatty liver disease; AID, autoimmune liver disease; HA, hepatic ascites; DM, Diabetes mellitus; HE, hepatic encephalopathy; VR, varicose vein rupture.

### Genotype counts and frequencies of the rs738409 G and rs58542926 T alleles in liver cirrhosis

Figure 1 shows genotype counts and frequencies of the rs738409 G allele. The frequencies of the CC, CG, and GG genotypes in rs738409 were 22.0%, 48.2%, and 29.8%, respectively. The highest frequency of the rs738409 G allele was 63.7% in the NAFLD. The lowest frequency of the rs738409 G allele was 45.5% in the virus. The frequencies of CC, CT, and TT in TM6SF2 rs58542926 were 81.7%, 16.1%, and 2.3%, respectively.

**Fig 1.**
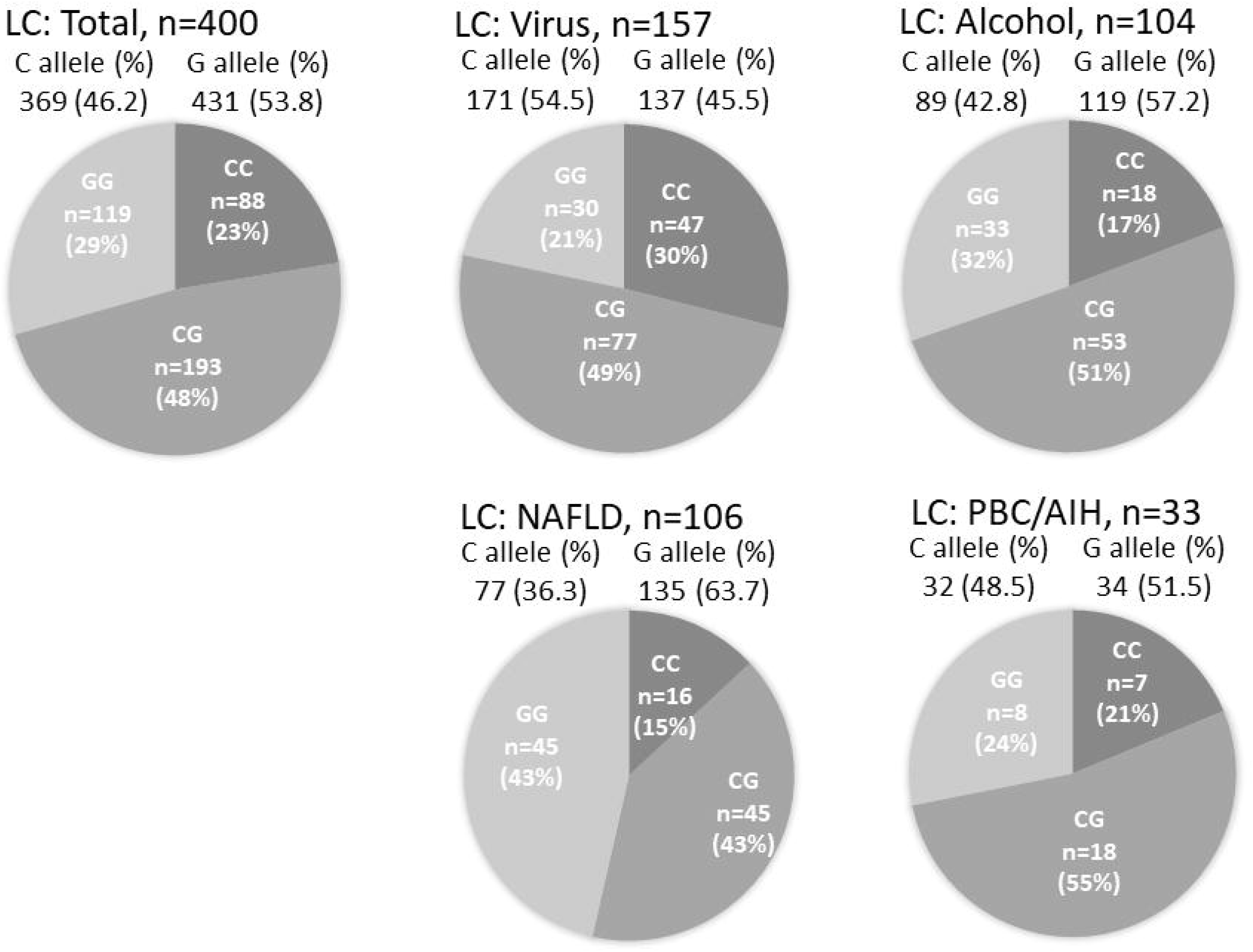
Genotype counts and frequencies of the rs738409 G allele in the LC group. LC, liver cirrhosis; NAFLD, nonalcoholic fatty liver disease.

Figure 2 shows genotype counts and frequencies of rs58542926 T allele. The highest frequency of the rs58542926 T allele was 16.0% in the NAFLD group of patients. The lowest frequency of the rs738409 T allele was 6.1% in the AIH. There was a significant difference between the frequencies of the mutated rs738409 and rs58542926 alleles (P < 0.001).

**Fig 2.**
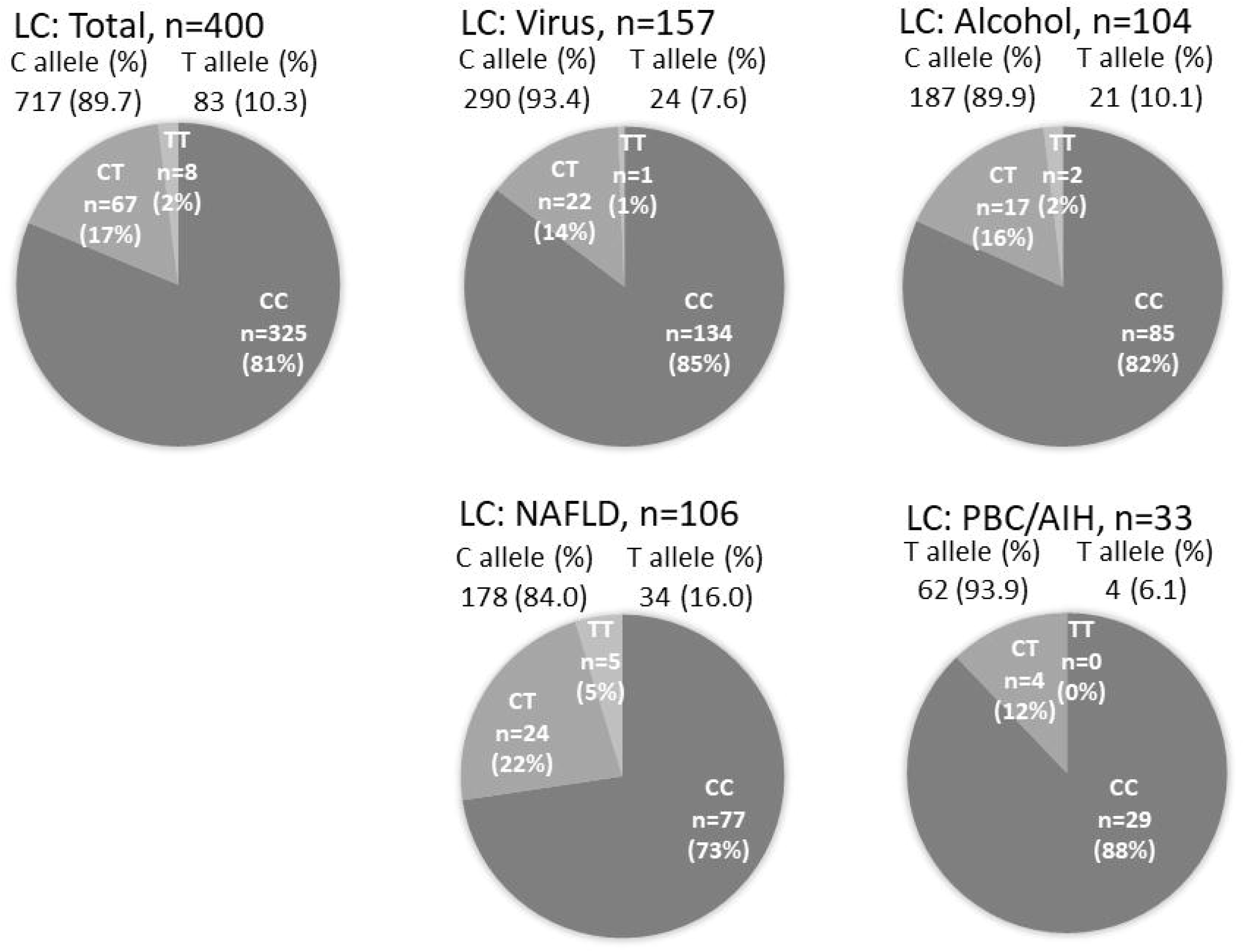
Genotype counts and frequencies of the rs58542926 T allele in the LC group. LC, liver cirrhosis; NAFLD, nonalcoholic fatty liver disease.

### Genotype counts and frequencies of the rs738409 G allele for cirrhosis-related complications

We evaluated the risk association between the G allele expression and complications in the patients (Table 2). In the NAFLD group, although there were no differences in frequencies of G alleles (OR = 2.260; 95% CI = 0.922–5.544; P = 0.075), the OR for complications significantly increased in the presence of the two mutated alleles (OR = 3.165; 95% CI = 1.073–10.294; P = 0.046) when the CC genotype was used as the reference. In the AID group, there was a significant risk of complications in the presence of the two mutated alleles (P = 0.014, respectively). On the other hand, there were no significant risks or complications in the virus and alcohol groups unless there was a presence of G alleles.

**Table 2.** Genotype counts and frequencies of PNPLA3 rs738409 for cirrhosis related complications

| Etiology | With complications<br>n (%) | Without complications<br>n (%) | OR | 95% CI | P value |
| --- | --- | --- | --- | --- | --- |
| Total | 199 (49.8) | 201 (50.3) |  |  |  |
|  | CC = 37 (18.6) | CC = 201 (50.3) | 1 | Reference |  |
|  | CG = 90 (45.2) | CG = 103 (51.2) | 1.204 | 0.724 - 2.004 | 0.474 |
|  | GG = 72 (36.2) | GG = 47 (23.4) | 2.112 | 1.205 - 3.699 | 0.009 |
|  | C allele = 164 (41.2) | C allele = 205 (51.0) | 1 | Reference |  |
|  | G allele = 234 (58.8) | G allele = 197 (49.0) | 1.401 | 0.916 - 2.144 | 0.120 |
| Virus | 75 (47.7) | 82 (52.3) |  |  |  |
|  | CC = 22 (29.3) | CC = 25 (30.5) | 1 | Reference |  |
|  | CG = 40 (53.3) | CG = 37 (45.1) | 1.229 | 0.594 - 2.541 | 0.579 |
|  | GG = 13 (17.3) | GG = 20 (24.4) | 0.739 | 0.299 - 1.823 | 0.511 |
|  | C allele = 84 (56) | C allele = 87 (53) | 1 | Reference |  |
|  | G allele = 66 (44) | G allele = 77 (47) | 0.904 | 0.458 - 1.786 | 0.771 |
| Alcohol | 57 (54.8) | 47 (45.2) |  |  |  |
|  | CC = 7 (12.3) | CC = 11 (23.4) | 1 | Reference |  |
|  | CG = 30 (52.6) | CG = 23 (48.9) | 2.050 | 0.688 - 6.110 | 0.198 |
|  | GG = 20 (35.1) | GG = 13 (27.7) | 2.418 | 0.745 - 7.845 | 0.142 |
|  | C allele = 44 (38.6) | C allele = 45 (47.9) | 1 | Reference |  |
|  | G allele = 70 (61.4) | G allele = 49 (52.1) | 1.371 | 0.592 - 3.176 | 0.046 |
| NAFLD | 51 (48.1) | 55 (51.9) |  |  |  |
|  | CC = 7 (13.7) | CC = 9 (16.4) | 1 | Reference |  |
|  | CG = 12 (23.5) | CG = 33 (60.0) | 0.468 | 0.142 - 1.534 | 0.210 |
|  | GG = 32 (62.7) | GG = 13 (23.6) | 3.165 | 1.073 - 10.29 | 0.046 |
|  | C allele = 26 (25.5) | C allele = 51 (46.4) | 1 | Reference |  |
|  | G allele = 76 (74.5) | G allele = 59 (53.6) | 2.260 | 0.922 - 5.544 | 0.075 |
| AIH | 16(48.5) | 17(51.5) |  |  |  |
|  | CC = 1 (6.3) | CC = 6 (35.3) | 1 | Reference |  |
|  | CG = 8 (50) | CG = 10 (58.8) | 4.800 | 0.475-48.46 | 0.184 |
|  | GG = 7 (43.8) | GG = 1 (5.9) | 42.00 | 2.136-825.7 | 0.014 |
|  | C allele = 10 (31.3) | C allele = 22(64.7) | 1 | Reference |  |
|  | G allele = 22 (68.8) | G allele = 12 (35.3) | 3.088 | 0.673 - 14.17 | 0.144 |

### Genotype counts and frequencies of the rs58542926 T allele for cirrhosis-related complications

We evaluated the risk association between the T allele expression and complications in the patients (Table 3). However, there were no significant risks or complications in the virus, alcohol, and NAFLD groups unless there was a presence of the T allele. Even in the NAFLD group analysis, there were no differences in frequencies of the T allele (OR = 0.843; 95% CI = 0.263–2.699; P = 0.772).

**Table 3.** Genotype counts and frequencies of the rs58542926 T allele for cirrhosis related complications

| Etiology | With complications<br>n (%) | Without complications<br>n (%) | OR | 95% CI | P value |
| --- | --- | --- | --- | --- | --- |
| Total | 199 (49.8) | 201 (50.3) |  |  |  |
|  | CC = 164 (82.4) | CC = 161 (80.1) | 1 | Reference |  |
|  | CT = 31 (15.6) | CT = 36 (17.9) | 0.845 | 0.499–1.432 | 0.532 |
|  | TT = [U+FF14] (2.0) | TT = 4 (2.0) | 0.982 | 0.241–3.992 | 0.979 |
|  | C allele = 359 (90.2) | C allele = 358 (89.1) | 1 | Reference |  |
|  | T allele = 39 (9.8) | T allele = 44 (10.9) | 0.897 | 0.443 - 1.816 | 0.783 |
| Virus | 75 (47.7) | 82 (52.3) |  |  |  |
|  | CC = 66 (88.0) | CC = 68 (82.9) | 1 | Reference |  |
|  | CT = 8 (10.7) | CT = 14 (17.1) | 0.589 | 0.232–1.496 | 0.265 |
|  | TT = 1 (1.3) | TT = 0 | n.c. |  | - |
|  | C allele = 140 (93.3) | C allele = 150 (91.5) | 1 | Reference |  |
|  | T allele = 10 (6.7) | T allele = 14 (8.5) | 0.797 | 0.219 - 2.896 | 0.729 |
| Alcohol | 57 (54.8) | 47 (45.2) |  |  |  |
|  | CC = 46 (80.7) | CC = 39 (83.0) | 1 | Reference |  |
|  | CT = 9 (15.8) | CT = 8 (17.0) | 0.954 | 0.336 - 2.708 | 0.929 |
|  | TT = 2 (1.9) | TT = 0 | n.c. |  | - |
|  | C allele = 101 (88.6) | C allele = 86 (91.5) | 1 | Reference |  |
|  | T allele = 13 (11.4) | T allele = 8 (8.5) | 1.325 | 0.318 - 5.524 | 0.698 |
| NAFLD | 51 (48.1) | 55 (51.9) |  |  |  |
|  | CC = 37 (72.5) | CC = 40 (72.7) | 1 | Reference |  |
|  | CT = 13 (25.5) | CT = 11 (25.5) | 1.278 | 0.510–3.203 | 0.601 |
|  | TT = 1 (2.0) | TT = 4 (7.3) | 0.270 | 0.029–2.530 | 0.252 |
|  | C allele = 87 (85.3) | C allele = 91 (82.7) | 1 | Reference |  |
|  | T allele = 15 (14.7) | T allele = 19 (17.3) | 0.843 | 0.263 - 2.699 | 0.772 |
| AID | 16 (48.5) | 17 (51.5) |  |  |  |
|  | CC = 15 (93.8) | CC = 14 (82.4) | 1 | Reference |  |
|  | CT = 1 (6.3) | CT = 3 (17.6) | 0.311 | 0.029–3.353 | 0.336 |
|  | TT = 0 | TT = 0 | n.c. |  | - |
|  | C allele = 31 (96.9) | C allele = 31 (91.2) | 1 |  |  |
|  | T allele = 1 (3.1) | T allele = 3 (8.8) | 0.405 | 0.012-12.28 | 0.607 |

### Potential confounding factors for the rs738409 polymorphism

A correlation between the rs738409 polymorphism and baseline clinical characteristics in the LC group was observed (Table 4). The BMI in the CC+CG group was significantly lower than that in the GG group (P = 0.004). The platelet count was higher in the CG+CC group than that in the GG group (P = 0.007). Hemoglobin A1c was lower in the CG+CC group than that in the GG group (P = 0.019). However, there were no associations between the rs738409 polymorphism and other hepatic enzymes or the total cholesterol.

**Table 4.**
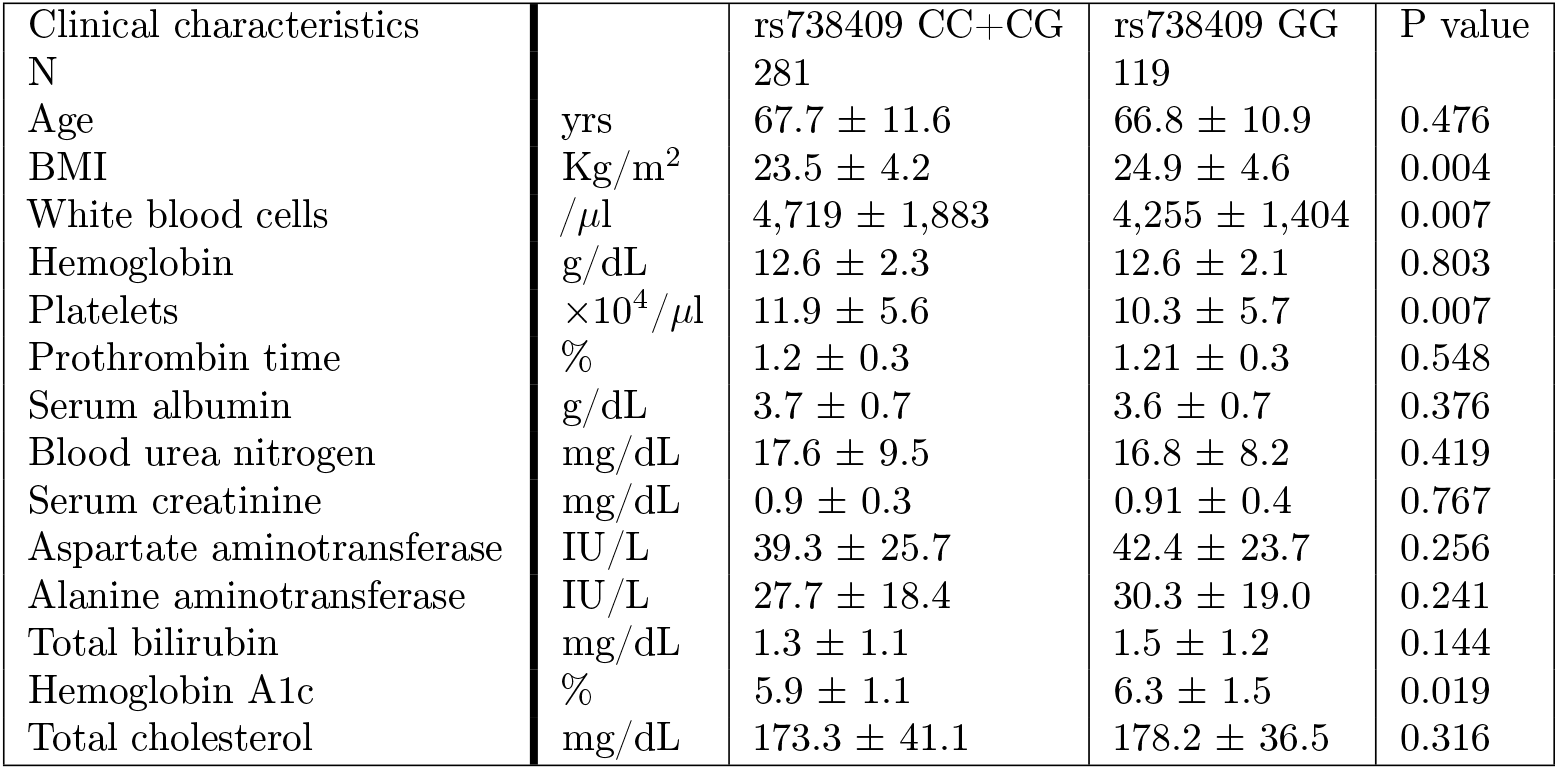
Effects of the PNPLA3 rs738409 polymorphism on clinical characteristics

| Clinical characteristics |  | rs738409 CC+CG | rs738409 GG | P value |
| --- | --- | --- | --- | --- |
| N |  | 281 | 119 |  |
| Age | yrs | 67.7 ± 11.6 | 66.8 ± 10.9 | 0.476 |
| BMI | Kg/m<sup>2</sup> | 23.5 ± 4.2 | 24.9 ± 4.6 | 0.004 |
| White blood cells | /μl | 4,719 ± 1,883 | 4,255 ± 1,404 | 0.007 |
| Hemoglobin | g/dL | 12.6 ± 2.3 | 12.6 ± 2.1 | 0.803 |
| Platelets | ×10<sup>4</sup>/μl | 11.9 ± 5.6 | 10.3 ± 5.7 | 0.007 |
| Prothrombin time | % | 1.2 ± 0.3 | 1.21 ± 0.3 | 0.548 |
| Serum albumin | g/dL | 3.7 ± 0.7 | 3.6 ± 0.7 | 0.376 |
| Blood urea nitrogen | mg/dL | 17.6 ± 9.5 | 16.8 ± 8.2 | 0.419 |
| Serum creatinine | mg/dL | 0.9 ± 0.3 | 0.91 ± 0.4 | 0.767 |
| Aspartate aminotransferase | IU/L | 39.3 ± 25.7 | 42.4 ± 23.7 | 0.256 |
| Alanine aminotransferase | IU/L | 27.7 ± 18.4 | 30.3 ± 19.0 | 0.241 |
| Total bilirubin | mg/dL | 1.3 ± 1.1 | 1.5 ± 1.2 | 0.144 |
| Hemoglobin A1c | % | 5.9 ± 1.1 | 6.3 ± 1.5 | 0.019 |
| Total cholesterol | mg/dL | 173.3 ± 41.1 | 178.2 ± 36.5 | 0.316 |

### Interaction between the rs738409 and rs58542926 polymorphisms

We investigated the interaction between the rs738409 and rs58542926 polymorphisms after adjusting for potential confounding factors (Table 5). In theNAFLD group, although there was no statistically interaction between the rs738409 and rs58542926 polymorphisms, the rs738409 GG + rs58542926 (CT+TT) group had the highest OR of 2.415 for cirrhosis related complications when rs738409 (CC+CG) + rs58542926 CC was used as the reference.

**Table 5.** Interaction between thePNPLA3 rs738409 and TM6SF2 rs58542926 polymorphism s in cirrhotic patients due to NAFLD

| Characteristic | Adjusted OR (95% CI) | P value |
| --- | --- | --- |
| rs738409 CC+CG + rs58542926 CC | 1 | Reference |
| rs738409 CC+CG + rs58542926 CT+TT | 0.657 (0.357–1.212) | 0.179 |
| rs738409 GG + rs58542926 CC | 1.500 (0.924–2.435) | 0.101 |
| rs738409 GG + rs58542926 CT+TT | 2.415 (0.886–6.582) | 0.085 |

## Discussion

To the best of our knowledge, this is the first study on the effects of the PNPLA3 and TM6SF2 polymorphisms on cirrhosis-related complications in Japanese patients with LC. PNPLA3 is a multifunctional enzyme and could regulate carbohydrate and lipid metabolism in the liver. This protein shows triglyceride hydrolase activity in hepatic cells and retinyl esters hydrolase activity in hepatic stellate cells. The polymorphism (rs738409 C>G p.I148M) in exon 3b of the PNPLA3 gene was reported to affect the levels of the PNPLA3 protein [20]. The cytosine (C) to guanine (G) substitution at this locus leads to reduced hydrolase activity and accumulation of large quantities of triglycerides in the liver [21]. As a result, the G allele of rs738409 has been strongly associated with advanced liver steatosis and hepatic inflammation [22].

The TM6SF2 protein is localized in the endoplasmic reticulum-Golgi intermediate compartment of human liver cells. The rs58542926 single nucleotide polymorphism (SNP) is characterized by C-to-T substitution in nucleotide 499, resulting in a glutamic acid to lysine substitution at codon 167 in the TM6SF2 gene (E167K) [23]. The rs58542926 polymorphism is likely to cause a loss of function of the TM6SF2 encoded protein and reduce the triglyceride secretion and increased intrahepatic triglyceride accumulation within multiple large lipid droplets in the liver tissue [24].

Three main results of the present study are most noteworthy. First, this study showed the frequency of the rs738409 G and rs58542926 T alleles in the Japanese people with LC. Especially, the rs738409 polymorphism had a clinical significance to evaluate the risk factor for the progression of fibrosis due to hepatic steatosis in LC patients.

Genotype counts and the frequency of the rs738409 G allele in the LC group were higher than those in the non-LC group regardless of etiology, which has also been previously reported [6–10]. The rs738409 polymorphism has been recognized as a risk factor for nonalcoholic steatohepatitis, which is characterized by the presence of steatosis leading to lipotoxicity and inflammatory damage to the major parenchymal cells in the liver [25–28]. Remarkably, the highest frequency of the mutated rs738409 allele in these genes was seen in the NAFLD group. On the other hand, its effect on disease progression in patients with hepatitis virus infection seems to be lower compared to its effect in patients with NAFLD.

Second, the PNPLA3 rs738409 GG genotype in the NAFLD group was a risk factor for the development of complications in the Japanese patients with LC. Some researchers had previously reported that G alleles of the rs738409 polymorphism significantly affected not only liver fat accumulation but also lead to increased susceptibility of developing severe hepatitis and a greater risk of fibrosis progression in non-LC patients [20]. Furthermore, several studies reported that the OR for the association between rs738409 and liver damage significantly increased in the GG genotype when the CC genotype was used as a reference [7, 21]. Patients with the rs738409 GG genotype need to be provided with the appropriate treatment to prevent fibrosis progression, particularly in the early stages of LC.

Third, the frequency of the rs58542926 T allele in the Japanese people with LC was relatively low, so that we could not accurately evaluate the association between rs58542926 and fibrosis progression. Ethnic differences in allele frequency of NASH (nonalcoholic steatohepatitis)-associated SNPs is well known [25]. This result was similar to those in two related previous studies which revealed a low frequency of the rs58542926 T allele in the population of northeast China [26, 27]. Regarding the rs58542926 T allele, there may have been scant clinical significance to adequately evaluate the progression of fibrosis among the Japanese people. However, the present study also revealed an additive effect of the rs738409 and rs58542926 polymorphisms in NAFLD for the Japanese patients with LC. Therefore, we recommend investigators to analyze the rs738409 and rs58542926 polymorphisms and, at the same time, to analyze the risk of cirrhosis related complications.

Genetic modifiers might be some of many factors that have strong interactions with the environment for the pathogenesis of triglyceride accumulation in hepatocytes. The pathogenesis of NAFLD involves complex interactions among environmental factors that lead to the development of severe obesity, metabolic syndrome, oxidative and endoplasmic reticulum stress, insulin resistance, lipotoxicity, and higher triglyceride levels [28]. In the present study, the frequencies of diabetes and obesity were higher in the rs738409 GG genotype than in the CC genotype in this cohort. However, the rs738409 polymorphism did not affect the levels of total cholesterol unless the risk alleles played a role in hepatic triglyceride hydrolysis. The reason for this discrepancy could be the severity of liver injury. The liver plays a central role in the secretion, degradation, synthesis, and storage of lipids. LC develops due to chronic liver damage slowly over months or years [29, 30]. As a result, total cholesterol levels tend to decrease with LC progression.

This study had a few limitations. First, this clinical study enrolled 400 patients with LC whose etiologies were not uniform. Especially, the number of patients with AID was relatively small. Although there were significant risks and complications identified in the AID group, further research is warranted. Second, the diagnosis of liver disease was based on laboratory results and imaging tests. Furthermore, the diagnosis of NAFLD did not meet the classical criteria for steatohepatitis because this diagnosis was not based on a liver biopsy in the LC patients. Finally, genomic DNA for this study was extracted from ethnically homogeneous Japanese people. Therefore, re-examination of the data for other and more diverse ethnic groups would be desirable.

In conclusion, to the best of our knowledge, this is the first study to determine the role of the PNPLA3 and TM6SF2 polymorphisms in fibrosis progression for Japanese patients with LC. The rs738409 GG genotype in the NAFLD group was a risk factor for LC-related complications. Furthermore, the additive effects of the rs738409 and rs58542926 polymorphisms had clinical significance to evaluate the risk factors for progression of fibrosis due to hepatic steatosis in patients with cirrhosis.

## Supporting information

available research data

## Acknowledgments

We thank the SATT Corporation for assistance with the statistical analysis. We also thank Robert E. Brandt, Founder, CEO, and CME, of MedEd Japan, for editing and formatting the manuscript.

## Author contributions

Xue Shao, Conceptualization, Writing Original Draft Preparation, & Review Haruki Uojima, Conceptualization, Writing Original Draft Preparation, Review & Editing

Taeang Arai, Yuji Ogawa, Toru Setsu, Masanori Atsukawa, Yoshihiro Furuichi, Yoshitaka Arase, Kazue Horio, Hisashi Hidaka, Takahide Nakazawa, Makoto Kako, Tatehiro Kagawa, Katsuhiko Iwakiri, Atsushi Nakajima, Shuji Terai, and Yasuhito Tanaka, Investigation

Wasaburo Koizumi, Supervision & Review

